# Self-regulation of the DNA N^6^-adenine methyltransferase AMT1 in the unicellular eukaryote *Tetrahymena thermophila*

**DOI:** 10.1101/2024.02.06.579081

**Authors:** Lili Duan, Haicheng Li, Aili Ju, Zhe Zhang, Junhua Niu, Yumiao Zhang, Jinghan Diao, Kensuke Kataoka, Honggang Ma, Ni Song, Shan Gao, Yuanyuan Wang

## Abstract

DNA N^6^-adenine methylation (6mA) is involved in gene transcription as a potential epigenetic mark in eukaryotes. Despite the reported methyltransferase (MTase) for 6mA methylation in several eukaryotes, the regulatory mechanisms that govern the activity of 6mA MTase remain elusive. Here, we exploited the 6mA MTase AMT1 to elucidate its self-regulation in the unicellular eukaryote *Tetrahymena thermophila*. Firstly, detailed endogenous localization of AMT1 was delineated both in vegetative and sexual stages, revealing a correlation between the 6mA reestablishment in the new MAC and the occurrence of zygotically expressed AMT1. Catalytically inactive AMT1 reduced 6mA level on the *AMT1* gene and its expression level, suggesting that AMT1 modulated its own transcription via 6mA. Furthermore, AMT1-dependent 6mA regulated the transcription of its target genes thus affecting the cell fitness, as demonstrated by manipulating the dosage of AMT1 using AMT1-RNAi strains. Our findings unveil a positive feedback loop of transcriptional activation on the *AMT1* gene and highlight the crucial role of AMT1-dependent 6mA for gene transcription.

## Introduction

DNA N^6^-adenine methylation (6mA) as a potential eukaryotic epigenetic mark is reported to be involved in transcriptional regulation in eukaryotes (1-15). However, the relationship of 6mA and transcription is diverse across the tree of life. Depending on the organisms, 6mA may either activate or repress transcription. For instance, 6mA is related with active transcription in the ciliate *Tetrahymena thermophila* (hereafter referred to as *Tetrahymena*), the unicellular alga *Chlamydomonas reinhardtii*, the nematode *Caenorhabditis elegans*, early-diverging fungi and the vascular plant *Arabidopsis thaliana* (2,4,5,10,11,16). In contrast, 6mA generally represses gene transcription in the lepidopteran *Bombyx mori*, the African clawed frog *Xenopus laevis*, zebrafish, and various mammalian cells (pig, mouse brain and human genome) (13,15,17-19). This divergence is most likely due to the differences in their 6mA methyltransferases (MTases). 6mA MTases have been identified in several eukaryotes, including N6AMT1 in human (19). DAMT-1 in *C. Elegans* (4), BmMETTL4 in *B. Mori* (18), METTL4 in mammalian mitochondrion (20), AMT1 in *Tetrahymena* (11), and MTA1 in *Oxytricha trifallax* (hereafter referred to as *Oxytricha*) (21). In *C. elegans, B. mori* and mammalian mitochondrion, their 6mA MTases, belonging to the METTL4 subclade of the MT-A70 family (22), are associated with low 6mA, non-consensus motifs and non-genic regions (4,18,20). In contrast, 6mA MTases found in *Tetrahymena* and *Oxytricha*, categorized as the AMT1 subclade of the MT-A70 family, are related with abundant 6mA, ApT motif, and Pol II transcription (11,21).

*Tetrahymena*, like most ciliates (23-25), possesses two types of nuclei, namely the transcriptionally active macronucleus (MAC) and the transcriptionally inert micronucleus (MIC) (23,26-32). In the vegetative stage, 6mA is exclusively present in the MAC, but not in the MIC (11). During the sexual stage (conjugation), the zygotic MIC differentiates into the new MAC for the sexual progeny, whereas 6mA in the developing new MAC needs to be re-established from the unmethylated germline MIC (11). We identified the major 6mA MTase AMT1 in *Tetrahymena* (11,33), but how AMT1 is regulated remains unknown.

In this study, we analyzed the cellular localization of AMT1 from different sources, including the parental MAC and zygotic MIC in *Tetrahymena*. We demonstrated that the timing of the reestablishment of 6mA in the new MAC correlated with that of zygotically expressed AMT1. Using a catalytically inactivated AMT1, we discovered that AMT1 modulated its own transcription by adjusting 6mA levels on its own gene. By manipulating the dosage of AMT1 in the AMT1-RNAi strain, we found AMT1-dependent 6mA regulated the transcription of its targeted genes, thus affecting the cell fitness of *Tetrahymena*.

## Materials and Methods

### Cell culture

Wild-type SB210 and CU428 strains of *Tetrahymena thermophila* were obtained from the *Tetrahymena* Stock Center (http://tetrahymena.vet.cornell.edu). Cells were cultured to the log phase (2-3×10 ^5^ cells/mL) in SPP medium at 30 °C (34). For mating, cells in the log phase were starved in 10 mM Tris-HCl, pH 7.4 for 16-18 h at 30 °C. Mating was initiated by mixing cells expressing different mating types (34). All strains were generated using the somatic and germline transformation with standard procedures (35,36), and detailed information were described in the supplemental materials and methods.

### Phenotypic analyses

For growth analyses, 0.8 × 10^5^ cells/mL cells were inoculated in 15 ml of SPP medium and incubated at 30°C. The cell concentration was measured at the indicated timepoints after inoculation by Coulter Counter Z2 (Beckman). For all strains, we measured concentrations in three biological replicates and performed statistical analysis as described (37).

### Immunofluorescence staining

Immunofluorescence staining was performed as previously described (11). The primary antibodies were incubated in a blocking buffer (1 × PBST, 10% Normal Goat Serum and 3% BSA) with the following dilution: 1:2000 in anti-6mA antibody (Synaptic Systems, 202003) and 1:200 in anti-HA antibody (Cell Signaling, #3724). These primary antibodies were detected by Goat anti-Rabbit IgG (H+L) - Alexa Fluor 555 (1:4000, Invitrogen, A-21428). For the co-staining, cells were incubated with anti-6mA antibody and anti-HA antibody (Covance, MMS-101P, 1:500 dilution), and these primary antibodies were detected by the secondary antibodies Goat anti-Rabbit IgG (H+L) - Alexa Fluor 555 and Goat anti-Mouse IgG (H+L) - Alexa Fluor 488 (Invitrogen, A11001, 1:4000), respectively. Zeiss Axio Imager Z2 and Olympus BX43 microscope were used for imaging.

### UHPLC-QQQ-MS/MS analysis

Genomic DNA was extracted by phenol-chloroform extraction as described previously (2). The DNA (500 ng) was incubated with DNase I (1U, NEB, M0303L), Fast AP (1 U, Invitrogen, EF0651) and snake venom phosphodiesterase I (0.005 U, Sigma, P4506) at 37 °C for 17 h, and digested into mononucleotides. The molecular masses of the mononucleotides were analyzed by Ultra-high-performance liquid chromatography-tandem mass spectrometry (UHPLC-QQQ-MS/MS) as described previously (10). The selective multiple reaction monitoring (MRM) transitions were detected under m/z 266/150 for 6mA and m/z 252/136 for dA. The ratio of 6mA/A was quantified by the calibration curves from nucleotide standards running simultaneously.

### Western blot

The cells were pelleted by trichloroacetic acid and lysed in the SDS sample buffer as described (38), and subjected to western blotting. Proteins on the blot were detected by anti-HA antibody (Cell Signaling, #3724, 1:2000) or anti-alpha-tubulin antibody (Sigma, T6199, 1:2000), followed by the incubation with the secondary antibodies Goat anti Rabbit IgG (H+L) HRP Conjugate (TransGen Biotech, HS101-01, 1:8000) and Goat Anti Mouse IgG (H+L) HRP Conjugate (TransGen Biotech, HS201-01, 1:8000), respectively.

### qPCR of 6mA IP samples

6mA immunoprecipitation (6mA IP) was performed as previously described (2), and detailed procedure was described in the supplemental materials and methods. For 6mA IP samples, we analyzed the enrichment of representative 6mA-methylated sites identified by SMRT-seq by qPCR using primers indicated in Table S1. Fold enrichment was calculated using the input DNA. The degree of 6mA enrichment was calculated by qPCR with the flanking primers to compare Ct values of methylated genes with that of unmethylated rDNA.

### 6mA IP-seq analysis

Sequencing adaptors of 6mA IP-seq (sequencing of 6mA IP samples) were trimmed, and short and low-quality reads were filtered out using Trim-Galore-0.6.7 (https://www.bioinformatics.babraham.ac.uk/projects/trim_galore/). The retained reads were mapped to the *T. thermophila* MAC reference genome (39) by Tophat-2.1.1 (40). Unique mapping reads were extracted in the mapping sam files. Then Picard MarkDuplicates-2.18.29 (https://broadinstitute.github.io/picard/) was used to remove redundant reads caused by PCR amplification. Enrichment of a certain feature by 6mA IP was defined as “(counts of IP reads in a certain feature/counts of total IP reads)/ (counts of input reads in a certain feature/counts of total input reads)”. 6mA enrichment in 6mA IP-seq was defined as regions where the enrichment ratio of IP to input was greater than 2. Normalized bigWig files and compared bigWig files were generated by Deeptools-3.5.1 to limit fragment lengths between 100-500 bp (41). IGV was used for visualization (https://software.broadinstitute.org/software/igv/). Reliable 300 bp bins from the groups 3, 4 and 5 were selected in a genome-wide scale. After the normalization for the counting numbers, these reads which aligned to the selected regions were then used to analyze 6mA enrichment in both gene regions and intergenic regions. To evaluate the relationship between 6mA and genes, 6mA occupancy was calculated using the 6mA penetration (the ratio of the methylated adenine numbers to the total numbers of adenine at this position) to represent the methylation level of a specific ApT site. 6mA occupancy in a certain gene was defined as “100 × the numbers of high-methylated 6mA site (penetration ≥ 0.8) in the gene/the gene length”, representing the numbers of high-methylated 6mA site per 100 bp in the gene. The high-methylated 6mA sites was obtained from the published SMRT-seq data (9).

### RNA sequencing

Total RNA from vegetatively growing WT and AMT1-RNAi cells treated with 1 μg/mL Cd^2+^ for 17 h, untreated AMT1-APPA, Δ*AMT1* and WT cells (∼2-3 × 10^5^ cells/mL), were extracted using Trizol reagent (Invitrogen, 15596026) as described previously (11). Sequencing libraries were generated using NEBNext Ultra RNA Library Prep Kit for Illumina (NEB, USA, Catalog #: E7530L) following the manufacturer’s recommendations. Index codes were added to the attribute sequences for each sample. RNA sequencing (RNA-seq) was performed by Illumina NovaSeq 6000. Trim Glore-0.6.7 was used to remove sequencing adapters and obtain high quality reads (length > 36 and quality > 20). Then retained reads were mapped to the *T. thermophila* MAC reference genome using Hisat2-2.1.0. Multiple mapping reads in the mapping sam file were deleted. After removing PCR duplicates using Picard MarkDuplicates-2.18.29, the retained reads were counted by FeatureCounts-1.6.1. Differentially expressed genes were calculated by DESeq2 in R (Padj < 0.05, log2FoldChange < -1 or > 1. Gene Ontology analysis was performed by TBtools-1-105.

### RT-qPCR

Total RNA was extracted using Trizol reagent (Invitrogen, 15596026), then reverse-transcribed using a HiScript III 1st Strand cDNA Synthesis Kit with random hexamer (Vazyme, R312-02). RT-qPCR was performed using EvaGreen Express 2 × qPCR MasterMix-Low ROX (Abm, MasterMix-LR) in 7500 Fast Real-Time PCR System as described previously (11). Two different pairs of primers were used for RT-qPCR of *AMT1* (AMT1-RTPCR-f4905/AMT1-RTPCR-r5155 and AMT1-qPCR-2415/AMT1-qPCR-2649) and *RAB46* (RAB46-f2018/ RAB46-r2246 and RAB46-f2091/RAB46-r2284). *JMJ1* (TTHERM 00185640, JMJ1-RTPCR-f2244/ JMJ1-RTPCR-r3176) were used as the internal control. All qPCR primers were listed in Table S1.

## Results

### Reestablishment of 6mA was correlated with the zygotically expressed AMT1

To investigate the regulatory mechanism of the 6mA MTase AMT1, we firstly analyzed its cellular localization. In our previous study, we analyzed the epitope-tagged AMT1 from an ectopic locus (OE-AMT1-NHA) (11). In this study, we generated cells expressing N-terminally hemagglutinin (HA) tagged AMT1 protein from its endogenous locus in the MAC (Supplementary Figure S1A-B, AMT1-NHA). In AMT1-NHA cells, 6mA level was comparable to that in wide-type (WT) cells (Figure 1A), confirming that the HA-tag did not interfere its methylation activity. Consistent with our previous observation in vegetative OE-AMT1-NHA cells (11), AMT1 localized exclusively in the MAC containing 6mA (Figure 1B and Supplementary Figure S1C).

**Fig. 1.**
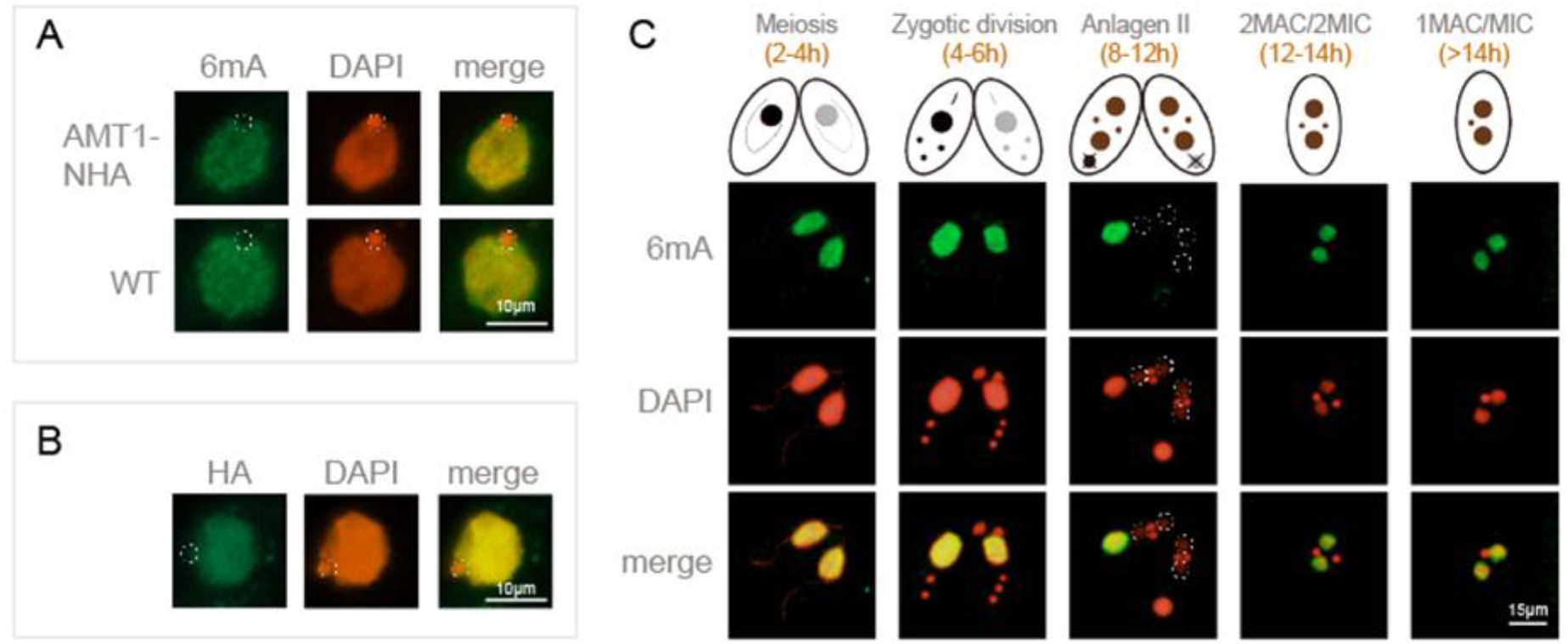
Cellular localization of AMT1 in the vegetative stage. **A.** Immunofluorescence (IF) staining using anti-6mA antibody (green) in AMT1-NHA and WT (SB210) cells. DNA was stained with DAPI (red). MIC was encircled by dotted line. **B.** IF staining of HA-tagged AMT1 in somatic AMT1-NHA cells. AMT1 was absent in the MIC (dotted circles). **C.** IF staining of 6mA in WT (SB210) cells during conjugation. 6mA was absent in the early new MAC (dotted circles).

During conjugation, the zygotic MIC differentiates into the new MAC (30), in which 6mA MTase(s) should be recruited to deposit 6mA *de novo* (42,43). By using AMT1-NHA-homozygous homokaryon (AMT1-NHA-homo/homo) cells, which expressed HA-tagged AMT1 from both the parental MAC and the zygotic MIC, we observed that AMT1 initially localized in the parental MAC and subsequently appeared in the new MAC soon after new MAC enlargement (Figure 2A, Anlagen II, Figure 2B, left panel). Intriguingly, 6mA was absent in new MAC at this stage (Figure 1C Anlagen II, Figure 2A top panel), which occurred approximately 4 hours after the new MAC formation (11) (Figure 1C, 2MAC/2MIC, Figure 2A top panel).

**Fig. 2.**
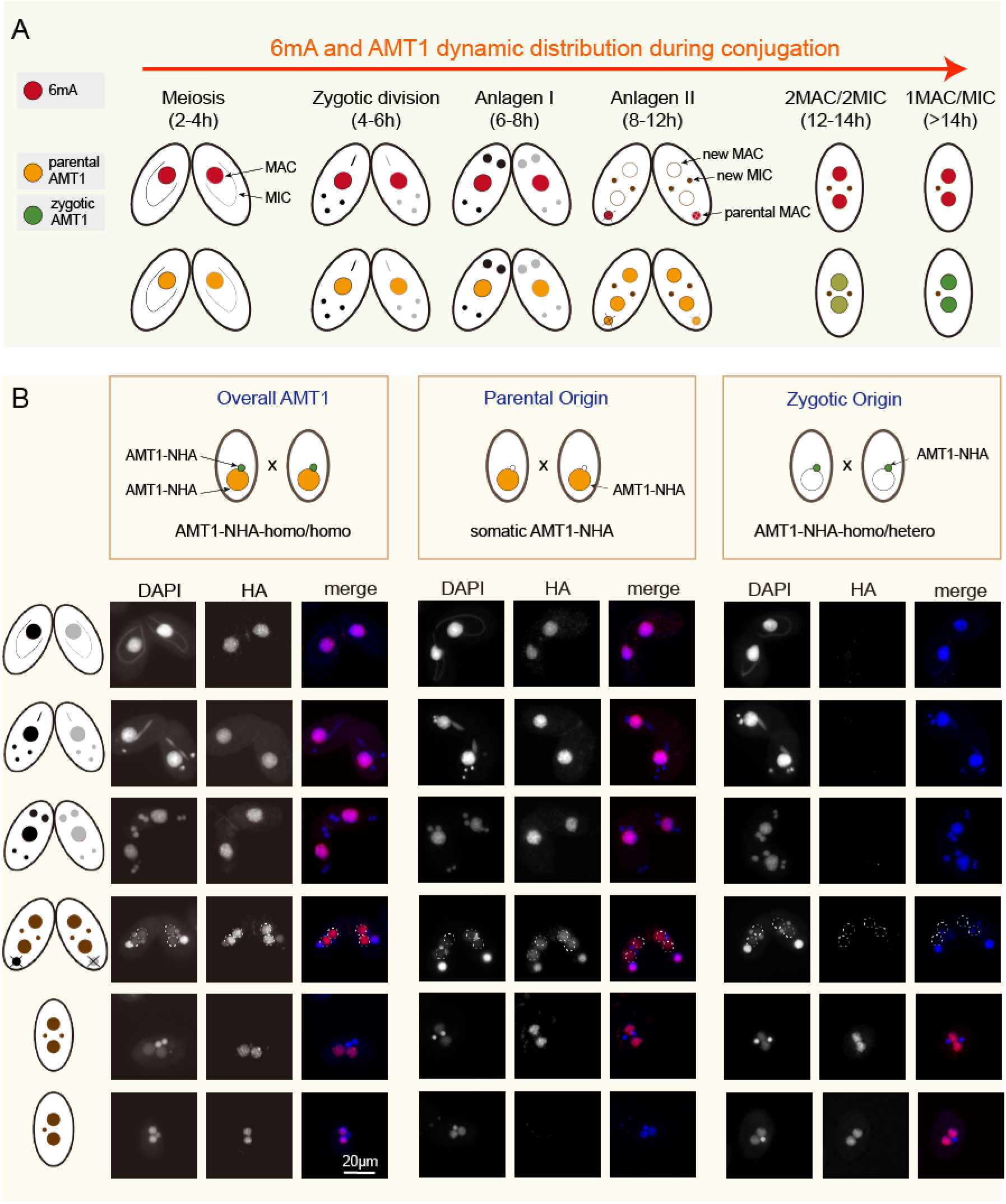
Cellular distribution of AMT1 during conjugation. **A.** Diagram showing the cellular localization of 6mA (top panel) and AMT1 (bottom panel) during conjugation. Nuclear events were used to ascertain conjugation stages. **B.** IF staining of HA-tagged AMT1 in AMT1-NHA-homo/homo, AMT1-NHA-homo/hetero and somatic AMT1-NHA strains during conjugation. Zygotic AMT1 was absent in the new MAC at the stage of Anlagen II (dotted circles), as demonstrated in AMT1-NHA-homo/hetero cells. Somatic AMT1 could last until the stage of 2MAC/2MIC.

To explore whether this delay of 6mA deposition in the early new MAC attributed to the lack of AMT1 activity, we generated strains expressing HA-tagged AMT1 either from the parental MAC (AMT1-NHA) or from the zygotic MIC (AMT1-NHA-homo/hetero) (Figure 2B). Somatic AMT1 first localized in the parental MAC during early conjugation and then in the new MAC lasting until the 2MAC/2MIC stage (Figure 2B, middle panel). In contrast, zygotic AMT1 started to appear in the new MAC at 2MAC/2MIC stage (Figure 2B, right panel), corresponding with the timing of 6mA.

### Self-regulation of AMT1 by AMT1-catalyzed 6mA

The MTase activity of AMT1 depends on the evolutionary conserved DPPW motif in the MT-A70 domain (Figure 3A). We previously discovered that the substitution of the DPPW motif to APPA for *AMT1* gene reduced the *AMT1* mRNA level, as demonstrated by RT-qPCR (11,22). To analyze this downregulation at the protein level, we generated cells expressing N-terminally HA-tagged AMT1 carrying the APPA mutation (AMT1-APPA-NHA) (Supplementary Figure S2A-B). Complete replacement of the AMT1 locus with the HA-tagged construct in the MAC was confirmed by Southern blot (Supplementary Figure S2C). Immunofluorescent (IF) staining showed that the AMT1 protein level was greatly reduced in AMT1-APPA-NHA cells compared with that in AMT1-NHA cells (Figure 3B and Supplementary Figure S2D). This reduction was confirmed by the Western blot analysis (Figure 3C).

**Fig. 3.**
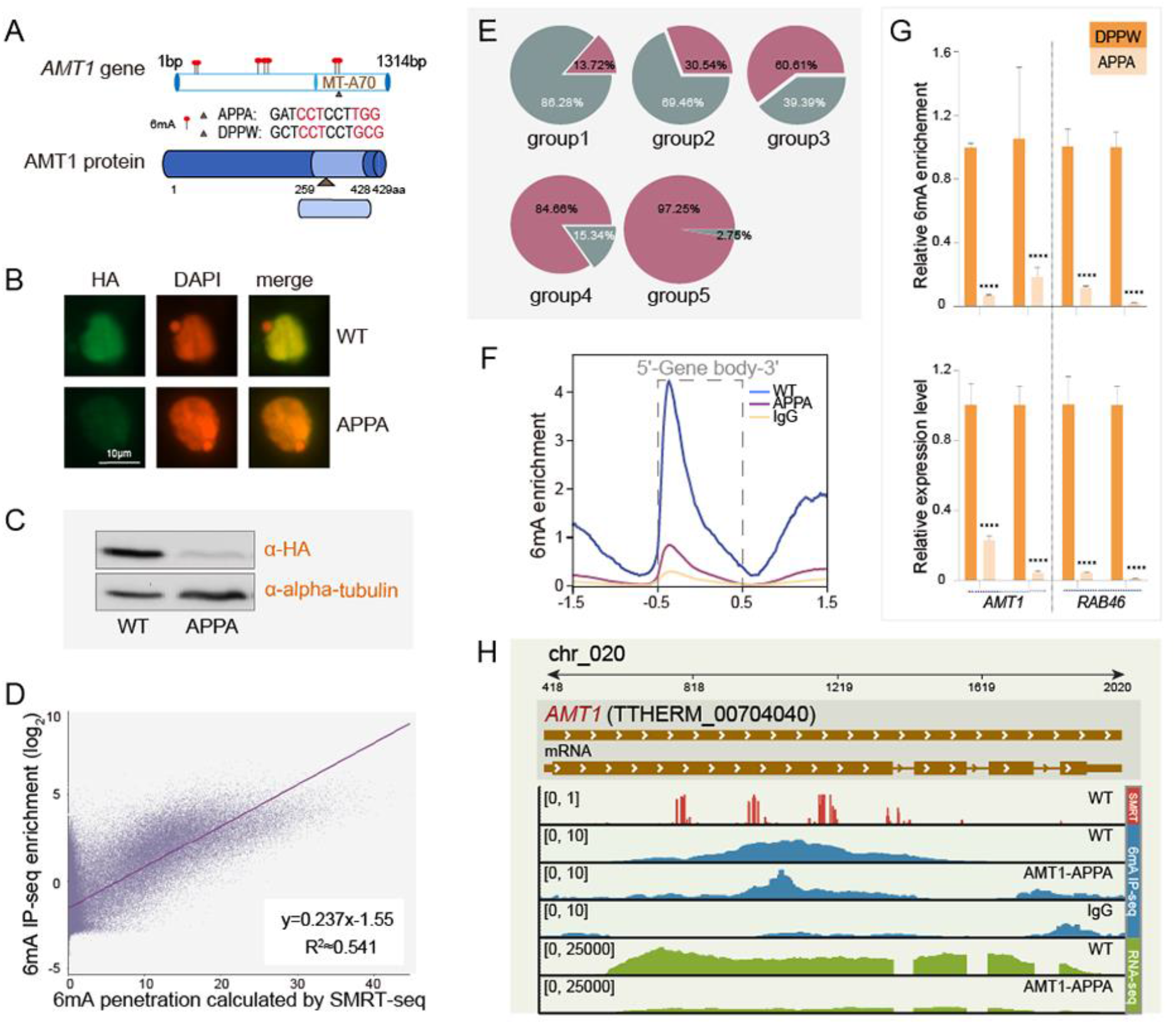
AMT1 regulated self-expression by 6mA. **A.** Illustration of *AMT1* gene locus and its domain structure. **B.** IF staining of HA-tagged AMT1 in AMT1-NHA (WT) and AMT1-APPA-NHA cells. **C.** Western blot of HA-tagged AMT1 in AMT1-NHA (WT) and AMT1-APPA-NHA cells. **D.** Correlation analysis of 6mA IP-seq and SMRT-seq in WT cells. 6mA penetration of a certain position was defined as “counts of 6mA/counts of total adenines” in SMRT-seq. The genome of *Tetrahymena thermophila* was split into 300bp regions, and the sum of penetration of all positions in a certain region was calculated to represent the 6mA methylation level of this region (Σ*P*). Enrichment of a certain region from 6mA IP-seq was calculated as “log_2_((counts of IP reads in the region/counts of total IP reads)/ (counts of input reads in the region/counts of total input reads))”. **E.** Evaluation of 6mA IP-seq enrichment for the regions with various 6mA methylation levels using SMRT-seq. Regions with the Σ*P* above 1.0 were divided into five groups according to 6mA methylation level from SMRT-seq data. From group 1 to group 5, the percentage of regions enriched by 6mA IP-seq were about 13.72%,30.54%, 60.61%, 84.66% and 97.25%. Note that the enrichment of 6mA IP-seq had very high coincidence with that of SMRT-seq in the regions with the high 6mA methylation level (*P* < 0.05). **F.** Distribution of 6mA peaks around the gene bodies in WT (blue) and AMT1-APPA cells (red) detected by 6mA IP-seq. IgG (orange) was used as the internal control. 6mA was accumulated downstream of TSS towards the 5’end of the gene body in WT and AMT1-APPA cells, but not in the IgG control. 6mA enrichment was defined as the ratio of normalized reads count between IP and Input. Note that only reads aligned into group 3, 4 and 5 regions were used to the above analysis. **G.** qPCR analysis of the 6mA IP samples and RT-qPCR analysis of mRNA showed that both 6mA levels and expression levels of *AMT1* and *RAB46* genes were more reduced in AMT1-APPA cells than that in WT cells, using two independent pairs of primers for each gene. Primers for unmethylated rDNA genes were used as the internal control for qPCR of the 6mA IP sample, and primers for the *JMJ1* gene were as the internal control for RT-qPCR of mRNA (Table S1). **H.** IGV snapshot of the *AMT1* gene locus. Tracks from top to bottom were: gene model, mRNA transcript, 6mA (WT cells using SMRT-seq, WT using 6mA IP-seq and AMT1-APPA cells using 6mA IP-seq), and RNA-seq coverage in WT and AMT1-APPA cells. Note that both 6mA level and *AMT1* expression were reduced in AMT1-APPA cells.

The decrease in AMT1 protein levels mirrored the observed reduction in *AMT1* mRNA levels.

Consistently, the APPA mutation reduced the global 6mA level in vegetative cells (Supplementary Figure S2E), and showed the similar phenotype with the Δ*AMT1* cells including slow cell growth and an abnormal contractile vacuole (Supplementary Figure S2F) (11,22). Furthermore, we found that 6mA was enriched on the *AMT1* gene body (1.37 6mA sites/100 bp, compared with 0.64 6mA sites/100 bp for other genes on average) (Figure 3A), raising the possibility that *AMT1* expression might be modulated by 6mA.

To address this possibility, we analyzed 6mA abundance on the *AMT1* gene in cells expressing the catalytically inactive AMT1 (AMT1-APPA) by 6mA-immunoprecipitation followed by DNA sequencing (6mA IP-seq). We first tested the reliability of 6mA IP-seq in WT cells. In WT cells, the 6mA IP-seq reads were enriched at the 5’ end of the gene body (Supplementary Figure S3A, green line). This enrichment was consistent with previously reported SMRT sequencing (SMRT-seq) analyses (9,21) (Supplementary Figure S3A, blue line). We then divided the *Tetrahymena* genome into 300 bp bins and compared the 6mA levels detected by SMRT-seq and by 6mA IP-seq. The results showed that most 6mA enriched regions found in 6mA IP-seq were also found in SMRT-seq (Supplementary Figure S3B). This correlation was more prominent along with the increase of 6mA methylation levels (R^2^≈0.541) (Figure 3D). Similarly, based on SMRT-seq, we categorized all bins into five groups according to their 6mA methylation levels (groups 1-5 correspond to the increased 6mA levels from low to high) (Figure 3E). Bins with high 6mA levels (97.25% of group 5, 84.66% of group 4 and 60.61% of group 3) were indeed accumulated with the reads from 6mA IP-seq (Figure 3E). These results indicated that 6mA IP-seq could reliably detect genomic regions enriched with 6mA especially for high 6mA regions (groups 3, 4, 5).

Next, we performed 6mA IP-seq for AMT1-APPA cells mainly focusing on the variation of high 6mA regions (groups 3, 4, 5). In comparison to WT cells, 6mA level of *AMT1* gene in AMT1-APPA cells was largely reduced (Figure 3G-H). Particularly, its 6mA enrichment at the 5’ end of gene body showed the significantly reduction (Figure 3F). Concomitantly, the *AMT1* mRNA was reduced in the AMT1-APPA cells (Figure 3G-H). Altogether, these results suggested that AMT1 promoted its own transcription by accumulating 6mA on its gene body.

The APPA mutation resulted in the appearance of abnormal contractile vacuole. This could be partially explained by the reduced expression of the Rab family GTPase *RAB46*, which plays an important role in contractile vacuole function in *Tetrahymena* (11). In line with our observation on the *AMT1* gene, the inactivation of AMT1 in AMT1-APPA cells affected both the 6mA and mRNA levels of the *RAB46* gene (Supplementary Figure S3C). This reduction was also confirmed by qPCR analysis of 6mA IP samples (Figure 3G and Table S1). These data suggested that the deactivation of AMT1 MTase activity led to the 6mA level reduction on the *RAB46* gene and thus diminished *RAB46* transcription.

### AMT1-dependent 6mA was involved in transcriptional regulation

In our previous study, we identified 6,047 differentially expressed genes (DEGs) (3,205 upregulated and 2,842 downregulated genes) upon *AMT1* knockout (11). However, it is important to note that these DEGs in Δ*AMT1* cells could be caused by cumulative effects. To minimize the cumulative effects, we generated AMT1-RNAi cells that could conditionally downregulate AMT1 by Cd^2+^ induction (Supplementary Figure S4A-B). To optimize the Cd^2+^ induction, we treated AMT1-RNAi cells with different concentrations of Cd^2+^ (0, 1, 3, 6 μg/mL). Our results showed that treatment with more than 1 μg/mL Cd^2+^ severely inhibited cell growth (Supplementary Figure S4C). Consistently, it was also reported that the growth rate with 1 μg/mL Cd^2+^ was indistinguishable from that in Cd^2+^ free cultured medium (44). Therefore, we decided to use 1 μg/mL of Cd^2+^ for subsequent experiments.

To trace the AMT1 protein level, we introduced the AMT1-NHA construct into the AMT1-RNAi cells (Supplementary Figure S4A-B). IF staining showed that the AMT1 protein level in the MAC of Cd^2+^ induced AMT1-RNAi cells gradually decreased over time (8 h, 11 h, 14 h, and 17 h) (Figure 4A and Supplementary Figure S4D). This gradual reduction was also confirmed by Western blot analyses (Figure 4B), indicating that AMT1 protein level could be reduced in inducible AMT1-RNAi cells.

**Fig. 4.**
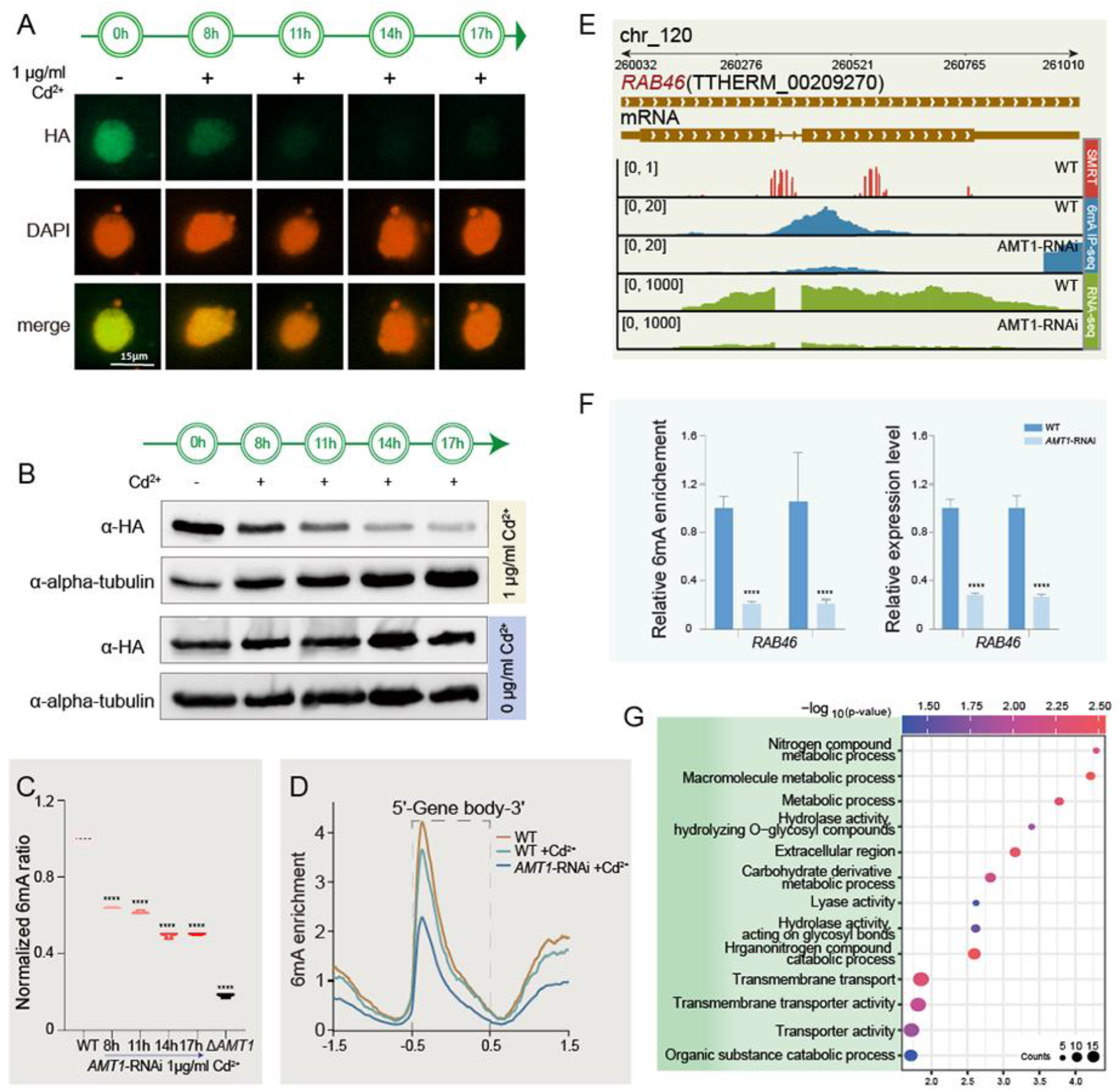
Gradual reduction of 6mA could be monitored by inducible AMT1-RNAi. **A.** IF staining of HA-tagged AMT1 in AMT1-RNAi cells induced by 1 μg/mL Cd^2+^ for 0 h, 8 h, 11 h, 14 h, and 17 h. **B.** Western blot of HA-tagged AMT1 in AMT1-RNAi cells induced by 1 μg/mL Cd^2+^ for 0 h, 8 h, 11 h, 14 h, and 17 h. AMT1-RNAi Cells without cadmium induction (0 μg/mL Cd^2+^) were collected at the corresponding timepoints for comparison. Alpha-tubulin was used as the loading control. **C.** Mass spectrometry analysis of 6mA in AMT1-RNAi cells induced by 1μg/mL Cd^2+^ for 8 h, 11 h, 14 h, and 17 h. WT and Δ*AMT1* cells were included as the controls to demonstrate the significant differences in 6mA ration. Three biological replicates were used for each sample. Data were presented as violin plots. Student’s t-test was performed. ^****^*P* < 0.001. **D.** Distribution of 6mA peaks around the gene body (TSS to TES) in uninduced WT (organe), induced WT (green) and induced AMT1-RNAi cells (blue) by 1 μg/mL Cd^2+^ for 17 h, detected by 6mA IP-seq. 6mA was accumulated downstream of TSS towards the 5’ end of the gene body in WT but declined in AMT1-RNAi cells. Note that only reads aligned into group 3, 4 and 5 regions were used to the above analysis. **E.** IGV snapshot of the *RAB46* gene locus (TTHERM 00209270). Tracks from top to bottom were: gene model, mRNA transcript, 6mA (WT cells using SMRT-seq, WT using 6mA IP-seq and AMT1-RNAi cells using 6mA IP-seq), and RNA-seq coverage in WT and AMT1-RNAi cells. Note that both 6mA level and *RAB46* expression were reduced in AMT1-RNAi cells. Both WT and AMT1-RNAi cells were induced with 1μg/mL Cd^2+^ for 17 h. **F.** qPCR analysis of the 6mA IP samples and RT-qPCR of mRNA showed that both 6mA levels and expression levels of *RAB46* genes were more reduced in AMT1-RNAi cells than that in WT cells, using two independent pairs of primers. Primers of unmethylated rDNA genes were used as the internal control for qPCR of the 6mA IP samples, and primers of the *JMJ1* gene were as the internal control for RT-qPCR of mRNA (Table S1). **G.** Gene ontology analysis for co-existing differentially downregulated genes (log_2_(Foldchange) < -1 and Padj < 0.05) with reduced 6mA level in AMT1-RNAi and Δ*AMT1* cells. The R package ggplot2 was performed. The size of dots represented the gene number. The color bar showed the *P*-value.

Next, we analyzed the 6mA level in AMT1-RNAi cells. MS analysis showed that 6mA level gradually declined with the prolonged time course in Cd^2+^ treated AMT1-RNAi cells (Figure 4C) but not in Cd^2+^ treated WT cells (Supplementary Figure S4E). Considering that the cells treated with Cd^2+^ for 17 hours dramatically reduced the global 6mA level (Figure 4C), we performed 6mA IP-seq in AMT1-RNAi cells under the same condition. We also mainly focused on the variation of high 6mA regions (groups 3, 4, 5) (Figure 3E). Although the 6mA enrichment in the 5’ end of gene body was still observed in AMT1-RNAi cells, its peak was largely reduced to about a half of WT cells (Figure 4D). This decline was not observed in Cd^2+^ treated WT cells (Figure 4D), suggesting that the diminishment of 6mA at the 5’ end of gene body in AMT1-RNAi cells was attributable to the reduction of AMT1.

We then compared the transcriptome between AMT1-RNAi and WT cells, both treated with 1 μg/mL Cd^2+^ for 17 hours. In total, 2,347 genes (9.1%) were upregulated and 1,134 genes (4.4%) downregulated in AMT1-RNAi cells (Supplementary Figure S4F). Considering 6mA is associated with active transcription in *Tetrahymena* (11), we focused on the downregulated genes. Among 1,134 downregulated genes, 544 genes exhibited both high enrichment of 6mA in WT cells and downregulated expression in AMT1-RNAi cells (Supplementary Figure S4G). We also conducted simultaneous RNA sequencing for Δ*AMT1* cells. About half of these genes (257 genes) displayed an overlap with genes that exhibited downregulation in both their 6mA and mRNA levels in Δ*AMT1* cells (Supplementary Figure S4G). Therefore, these 257 genes were considered credible as AMT1-regulated genes based on these analyses. Specifically, 6mA IP followed by qPCR and RT-qPCR showed that both the 6mA level and the mRNA level of *RAB46* were also significantly reduced in AMT1-RNAi cells (Figure 4E-F and Supplementary Figure S4F), suggesting *RAB46* was indeed regulated by AMT1-dpendent 6mA.

Gene ontology (GO) analysis of these 257 genes revealed the association with various biological pathways including macromolecule, nitrogen and carbohydrate derivative metabolic processes, organic substance catabolic processes and transmembrane transport (Figure 4G). The downregulation of genes that were involved in various metabolic pathway-related processes could explain the growth defect when AMT1 was impaired, and the abnormally large contractile vacuole might be linked with the dysfunction of transmembrane transport related processes. Therefore, AMT1-dependent 6mA regulated the transcription of these genes, thus affecting cell fitness of *Tetrahymena*.

## Discussion

As a novel DNA modification in eukaryotes, 6mA is reported to be regulated by its methyltransferases. However, the regulatory mechanism of 6mA MTase remains elusive. Here, we demonstrated both self-regulation and transcriptional regulation of 6mA MTase AMT1 in *Tetrahymena* (Figure 5).

**Fig. 5.**
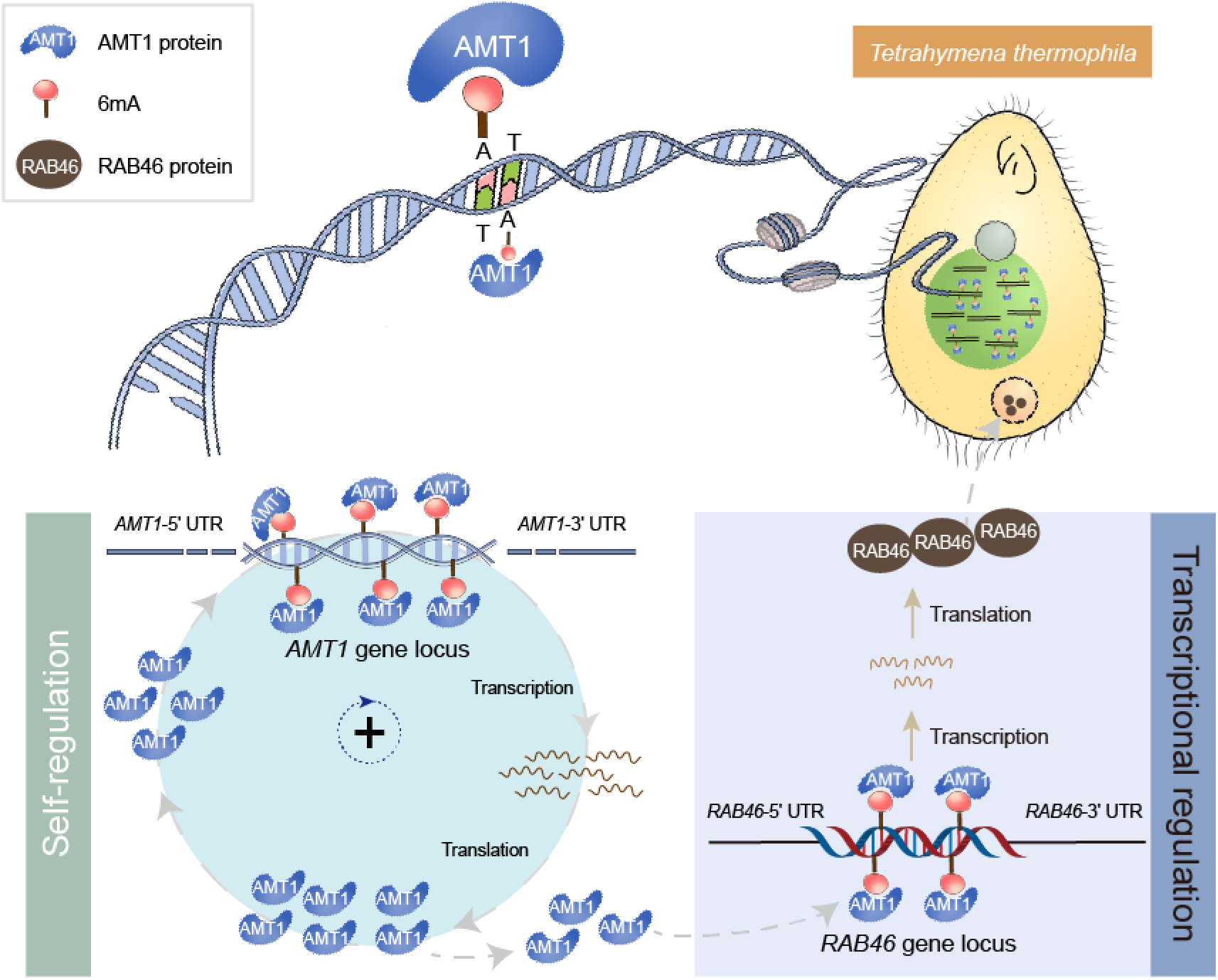
Diagram denoting AMT1 self-regulation and gene transcriptional regulation by AMT1-dependent 6mA.

The 6mA methyltransferase AMT1 was localized in the new MAC immediately after its formation (Figure 2A, Anlagen II), whereas 6mA only appeared in the new MAC approximately 4 hours after the new MAC formation (10,11) (Figure 2A, 2MAC/2MIC). In this study, we analyzed the localization of AMT1 protein from the parental MAC or the zygotic MIC. We found that that although AMT1 from the parental MAC existed in the new MAC at the stage of Anlagen II, 6mA did not appear in the same nucleus. Instead, the timing of 6mA appearance correlated with that of AMT1 expressed from the zygotic MIC at the stage of 2MAC/2MIC, suggesting that AMT1 in the early stage of new MAC was not capable of depositing 6mA. There are two scenarios that are not mutually exclusive. First, the catalytic activity of AMT1 requires cofactors that may localize to the new MAC during the stage of 2MAC/2MIC. In vegetative cells, the 6mA methyltransferase AMT1 (also known as MTA1) forms complex with MTA9, p1 and p2, and the methyltransferase activity of AMT1 is only detected in the presence of MTA9, p1 and p2 in vitro (21,45,46). In the new MAC,these cofactors may be limiting factors for the catalytic activity of the AMT1. Second, since the preferable substrates of AMT1 are hemi-methylated ApT dinucleotides (9), unmethylated ApT dinucleotides in the early new MAC may not be methylated by AMT1 until hemi-methylated 6mApT is prepared *de novo* possibly by other 6mA MTases. Further research is needed to address these hypotheses.

During the vegetative stage, 6mA levels do not change significantly even after DNA replication (10,11). This can probably be attributed to the high level of AMT1 activity (9). Self-regulation of AMT1 is an intelligent way to achieve effective and efficient methylation by the positive feedback, enhancing its methylation ability by methylating its own gene.

APPA mutation and RNAi of *AMT1* abolished its MTase activity and contributed to the decline of 6mA levels in the *AMT1* gene, which in turn regulated *AMT1* transcription. Similarly, when AMT1 was downregulated, the 6mA levels of *RAB46* gene also declined, thus contributing to its lower expression. These findings support the correlation of 6mA and transcription. The association between 6mA and global transcription turned out to be weak (10,11), which could be attributed to the transcriptional co-regulation by several factors including 6mA and various histone marks (e.g., H2A.Z, H3K4me3). When 6mA levels are decreased, other active marks could be upregulated to maintain the balance of gene transcription.

## Data availability

All data generated in this study can be accessed through BioProject accession number PRJNA1033614.

## Author contributions

YW and SG conceived the study, supervised most of the experiments and analyzed the data. LD performed most of the experiments. HL performed the bioinformatic analyses. ZZ generated HA tagged AMT1 strains with AJ. JN carried out the UHPLC-QQQ-MS/MS analyses with NS. YZ detected the expression level of *RAB46* gene. JD modified the language. KK designed and supervised the Southern blot performed by LD. YW, SG and KK prepared the manuscript with LD and HL. All authors read and approved the final manuscript.

## Acknowledgements

The authors would like to thank the following people for assistance with this study: Congmingmei Liu (Ocean University of China, OUC) for drawing the AMT1 model, Dahua Chen (Institute of Biomedical Research, Yunnan University) for providing the platform of UHPLC–QQQ–MS/MS and Wenxin Zhang (Institute of Biomedical Research, Yunnan University) for supervising the UHPLC-QQQ-MS/MS analysis, Alan Warren (Department of Life Sciences, Natural History Museum) for polishing the language. Our special thanks are given to Dr. Weibo Song (OUC) for his kind suggestions during preparation of the manuscript. High-performance computing resources for data processing were provide by the Institute of Evolution & Marine Biodiversity at OUC, the Center for High Performance Computing and System Simulation at Laoshan Laboratory, and Marine Big Data Center of Institute for Advanced Ocean Study at OUC.

